# Reparametrizing the Sigmoid Model of Gene Regulation for Bayesian Inference

**DOI:** 10.1101/352070

**Authors:** Martin Modrák

**Affiliations:** Institute of Microbiology of the Czech Academy of Sciences, Prague, Czech Republic

**Keywords:** Gene Regulation, Gene Network Inference, Bayesian Statis-tics

## Abstract

This poster describes a novel reparametrization of a frequently used non-linear ordinary differential equation (ODE) model of gene regulation. We show that in its commonly used form, the model cannot reliably distinguish between both quantitatively and qualitatively different parameter combinations. The proposed reparametrization makes inference over the model stable and amenable to fully Bayesian treatment with state of the art Hamiltonian Monte Carlo methods.

Complete source code and a more detailed explanation of the model is available at https://github.com/cas-bioinf/genexpi-stan.

## 1 Introduction

Transcriptional regulation has historically been treated with a vast range of models from Boolean networks, to full Michaelis-Menten simulation of individual chemical reactions. A middle ground is occupied by linear and non-linear ODE models that approximate Michaelis-Menten kinetics. In our attempts to reimplement a Bayesian version of a popular sigmoid ODE model [1] we noticed that very different parameter values - including different sign of the regulatory effect - may result in almost identical behavior, which introduces computational difficulties and questions the interpretation of previous results achieved with the model. Here we describe novel reparametrization of the model implemented in the Stan probabilistic programming language [2] that mitigates these problems.

## 2 The Model

The ODE model we use is based on [3]. For a bacterial regulon with a single regulator *y* and targets *x*_1_…*x*_*n*_ the model takes form

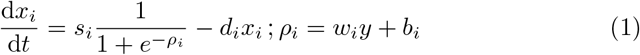

Where *s*_*i*_, *w*_*i*_, *b*_*i*_ and *d*_*i*_ are parameters to be fit. While bacterial regulons are our primary interest, the model also works with multiple regulators. The model assumes that both the regulator and targets are measured at multiple time points, letting us to solve the ODE. When using microarray data we assume normal observation noise where the standard deviation has two components: one constant and one proportional to the expression. The regulator is fit using B-splines where the spline coefficients are also treated as parameters of the model which let us to both handle uncertainty in regulator measurements and obtain the regulator at arbitrarily small time resolution required for solving the ODE numerically. We note that the model is only weakly identifiable in its direct form (1), as very different parameter sets can result in similar solutions for *x*_*i*_. Weak identifiability poses computational problems for Bayesian inference, reduces stability of maximum-likelihood estimates and limits interpretability of the model results. Critically, the sign of *w*_*i*_ (whether the regulator is an activator or a repressor) is not well determined for certain target profiles (Fig. 1a) and it may be impossible to determine whether *w*_*i*_ ≃ 0 (Fig. 1b). Negligible effect on *x*_*i*_ can also be observed under linear transformations of (*w*_*i*_, *b*_*i*_) when |*w*_*i*_*y*+*b*_*i*_| ⪢ 0 (Fig. 1c) and under approximately linear transformations of (*s*_*i*_,*d*_*i*_) (Fig. 1d).

**Fig. 1.**
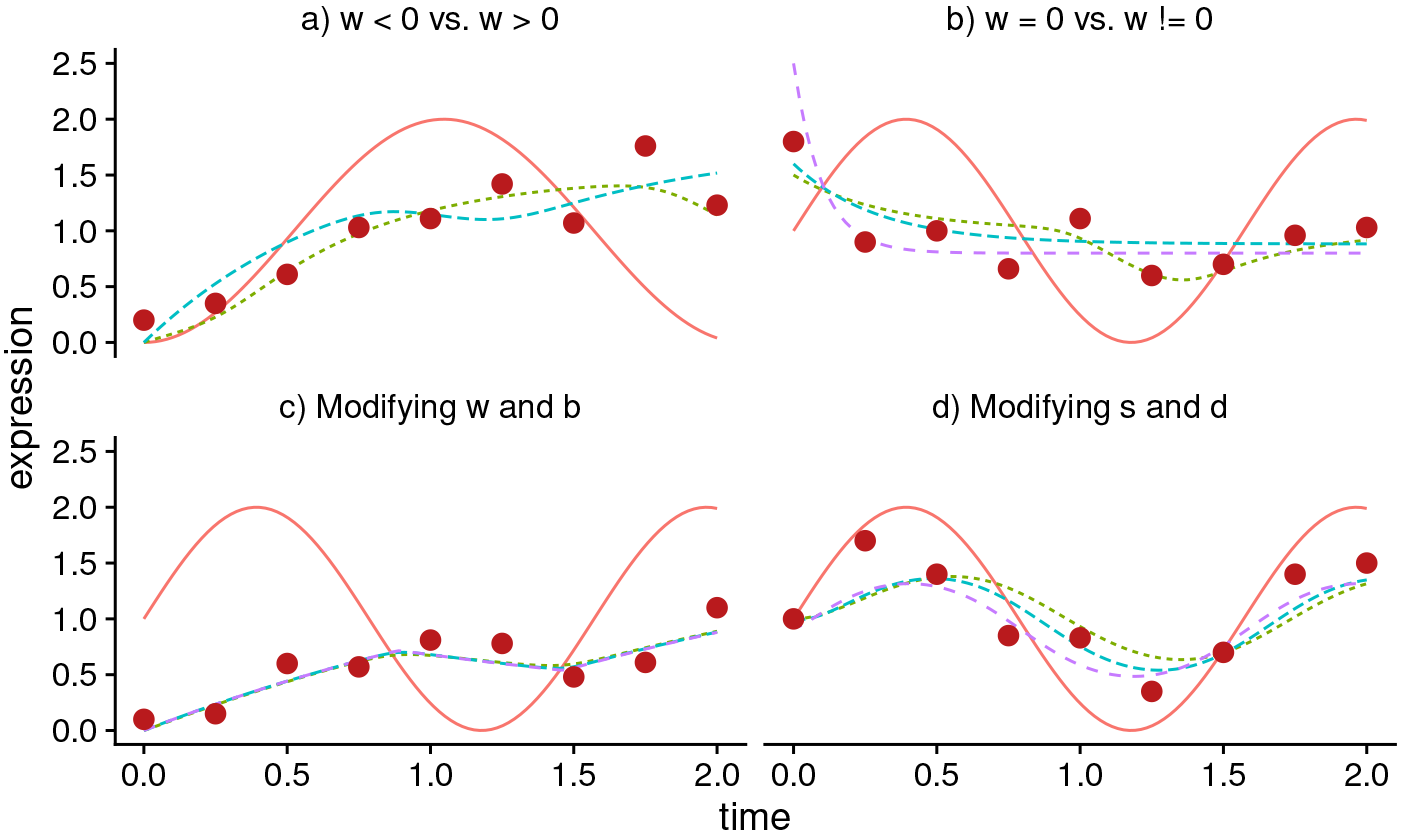
Simulated examples of weakly identified parameters in the sigmoid model. Expression of the regulator (red, solid line) and multiple significantly different parameter values that give raise to similar solutions of the ODE (dashed lines) are shown together with possible measured target profiles (dots) that are insufficient to distinguish between the solutions. a) (*w*,*b*,*s*,*d*) ∈ {(5, −1, 3, 2), (−5,10, 2.5,1.4)} b) (*w*,*b*,*s*,*d*) ∈ {5, −1, 3, 3), (0, 0, 6, 3.4), (0, 0,16,10) c) *s* = 1, 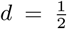 and (*w*,*b*) ∈ {(5, −3), (10, −6), (50, −30)} d) *w* = 1,*b* =-1and(*s*,*d*) ∈ {(10, 5), (19,10), (180,100)}

The aforementioned identifiability problems can be mitigated by a) fixing *I*_*i*_ = *sgn*(*w*_*i*_) and running separate fits when both signs have to be tested and b) replacing *w*_*i*_,*b*_*i*_ and *s*_*i*_ with 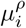, the mean regulatory input 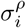, the std. deviation of the regulatory input and *a*_*i*_ - normalized ratio of *s*_*i*_ to *d*_*i*_. The original parameters can then be recovered as:

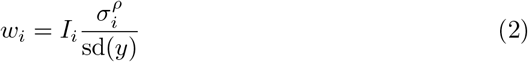

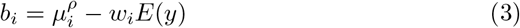

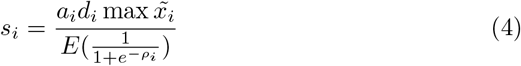

where *E* and sd correspond to the sample mean and standard deviation and 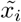 is the observed expression of gene *i*.

To our knowledge, this is the first time the identifiability issues are reported and handled for the sigmoid ODE model. Caution should therefore be exercised when interpreting the parameter values found in previous works using this model.

## 3 Workflow and Remarks

A model fit is in itself insufficient to determine whether a regulation is plausible. To assess plausibility, we test whether the model improves over two baselines: a) constant synthesis (*w*_*i*_ = 0) and b) a model where the regulator spline is not constrained by regulator measurements data. We use the LOO-IC criterion [4] to compare the models.

When there are known regulations, the model can be trained by fitting those known regulations first. The posterior estimates of the expression of the regulator can then be used to decrease uncertainty when fitting putative novel targets. Performance-wise, the training phase takes several minutes and fitting 881 putative targets individually on an 8-core machine takes 1 hour which promises faster performance than [1].

